# Perturbed sulfur homeostasis allows *C. elegans* to escape growth retardation on *Actinobacteria* from its natural microbiome

**DOI:** 10.1101/2023.11.12.566774

**Authors:** Om Patange, Peter Breen, Gary Ruvkun

## Abstract

The rate at which organisms grow is influenced by their biotic environment. The nematode *Caenorhabditis elegans* grows slower in the presence of *Actinobacteria*, but it is unknown why. Here, we show how perturbed levels of hydrogen sulfide and cysteine modulate the growth rate of *C. elegans* on *Actinobacteria*. Using an unbiased forward genetic screen of *C. elegans* we discovered alleles of the conserved cystathionine gamma-lyase (*cth-2/CTH)* that improved animal growth rate on *Actinobacteria*. Conversely, null alleles of *cth-2* cause developmental arrest of animals grown on *Actinobacteria*, which can be rescued by exogenous H_2_S. We also discovered a leucine rich repeat gene that regulates cysteine and H_2_S production, *lrr-2/LRRC58.* A wild isolate of *C. elegans* that naturally grows well on *Actinobacteria* has a mutant allele of *lrr-2*, suggesting this sulfur metabolism pathway is important for the regulation of animal growth rate in its natural ecological context. We propose a model in which wild-type worms use sulfurous compounds to promote growth of their favored bacterial food sources by inhibiting *Actinobacteria* growth. This strategy becomes a liability when *Actinobacteria* are the sole food source but can be bypassed by mutations in sulfur metabolism. This study reveals how the homeostasis of sulfurous compounds controls the growth rate of animals in an ecological context.

## Introduction

Many molecular mechanisms of how species interact have been discovered, such as innate and adaptive immune systems^1^, and community assembly mediated by metabolite exchange^2^. However, given a pair of interacting species, we cannot *a priori* predict the relevant molecules exchanged nor their ecological dynamics^3^. One reason for this is that a large fractions of genes in even the best studied organisms remain uncharacterized^4^. *Caenorhabditis elegans* has been a powerful tool to discover the molecular basis of diverse ecological interactions. For example, the pathogenesis of *Pseudomonas aeruginosa*^5,6^ and *Microbacterium nematophilum*^7^; the chemical modulation by bacteria of animal behavior^8,9^; the exchange of vitamins^10,11^, siderophores^12^ and cofactors^13^ between microbes and animals; and the role of metabolites^14^ and RNA^15^ in transgenerational epigenetic memory.

The wild ecology of *C. elegans* has recently been characterized, facilitating many of these discoveries^16–19^. In the wild, *C. elegans* grows on rotting vegetation containing a rich diversity of microbes including the dominant bacterial clades of *Proteobacteria, Firmicutes, Bacteroidetes,* and *Actinobacteria;* while in the lab it is usually fed a diet of the *Proteobacteria Escherichia coli*. The growth rate and fecundity of *C. elegans* depends strongly on the bacterial composition of its environment^16,17^. In particular, rotting vegetation with a high abundance of *Actinobacteria* has been found to not support *C. elegans* proliferation^17^. Furthermore, *C. elegans* produces a smaller brood when it is cultured on individual species of *Actinobacteria*^16^. However, it remains unclear what causes *C. elegans* to grow poorly on *Actinobacteria*.

Species of the *Actinobacteria* clade are found in diverse environments including rotting vegetation^17^, plant root microbiomes^20^, and human gut microbiomes^21^. The secondary metabolism of *Actinobacteria* is a major source of clinically relevant antibiotics^22^. *Mycobacterium tuberculosis*, the etiological agent of tuberculosis, is a member of the *Actinobacteria* clade. Sulfur metabolism, in particular the exchange of hydrogen sulfide (H_2_S), has been found to mediate the interaction of animals and *M. tuberculosis*^23–25^. The trans-sulfuration pathway that generates cysteine from homocysteine has been found to play an important role in mouse models of tuberculosis. When either *CBS*, the enzyme that metabolizes homocysteine to cystathionine, or *CTH*, the enzyme that metabolizes cystathionine to cysteine are mutated in mice, they are better able to survive against *M. tuberculosis* infection^23,24^. Both of these enzymes have also been shown to catalyze reactions that make H_2_S^26^. Hydrogen sulfide, in turn, has been found to improve the resistance of various bacteria to antibiotics^27^ as well as improve the growth of *M. tuberculosis*^25^. H_2_S is also emerging as an endogenously protective gas for animals, for instance it has been found to increase *C. elegans* longevity^28^. Indeed, *C. elegans* maintains sulfur homeostasis in part by sensing H_2_S levels^29,30^, suggesting it is an important metabolite for the animals.

Here, we used an unbiased genetic approach to discover the molecular basis of growth retardation of *C. elegans* cultivated solely on *Actinobacteria*. Our genetic screen revealed the importance of the trans-sulfuration pathway in modulating animal growth rate. In contrast to the case of animals infected with *M. tuberculosis*, wild-type *(WT) C. elegans* produces excess H_2_S and this is deleterious to *Actinobacteria* in its environment. Reducing H_2_S with mutations in the sulfur metabolism pathway improves growth of animals on *Actinobacteria,* but abolishing H_2_S production renders animals unable to grow on *Actinobacteria*. We find lesions in sulfur metabolism genes in wild isolates of *C. elegans* suggesting modulation of H_2_S is ecologically important.

## Results

### 1. Regulation of sulfur amino acids is key to development of *C. elegans* on *Actinobacteria*

The population of *Actinobacteria* in the natural microbiome of *C. elegans* increases from ∼1% to ∼7% of the total bacterial population when these animals fail to grow well^17^. To assay the effect of *Actinobacteria* on *C. elegans* growth, we fed animals a diet of only *E. coli* OP50 or only *Microbacterium* JUb115. *E. coli* is the standard *C. elegans* lab diet and a *Proteobacteria,* the dominant clade in the *C. elegans* microbiome; while *Microbacterium* JUb115 is an *Actinobacteria* isolate from the *C. elegans* microbiome^17^. We found a stark delay in development (Figure 1A,B). Animals grew to egg-laying adulthood ∼60% more slowly (Figure 1B), and produced a smaller brood by ∼10-fold (Figure 1F).

**Figure 1:**
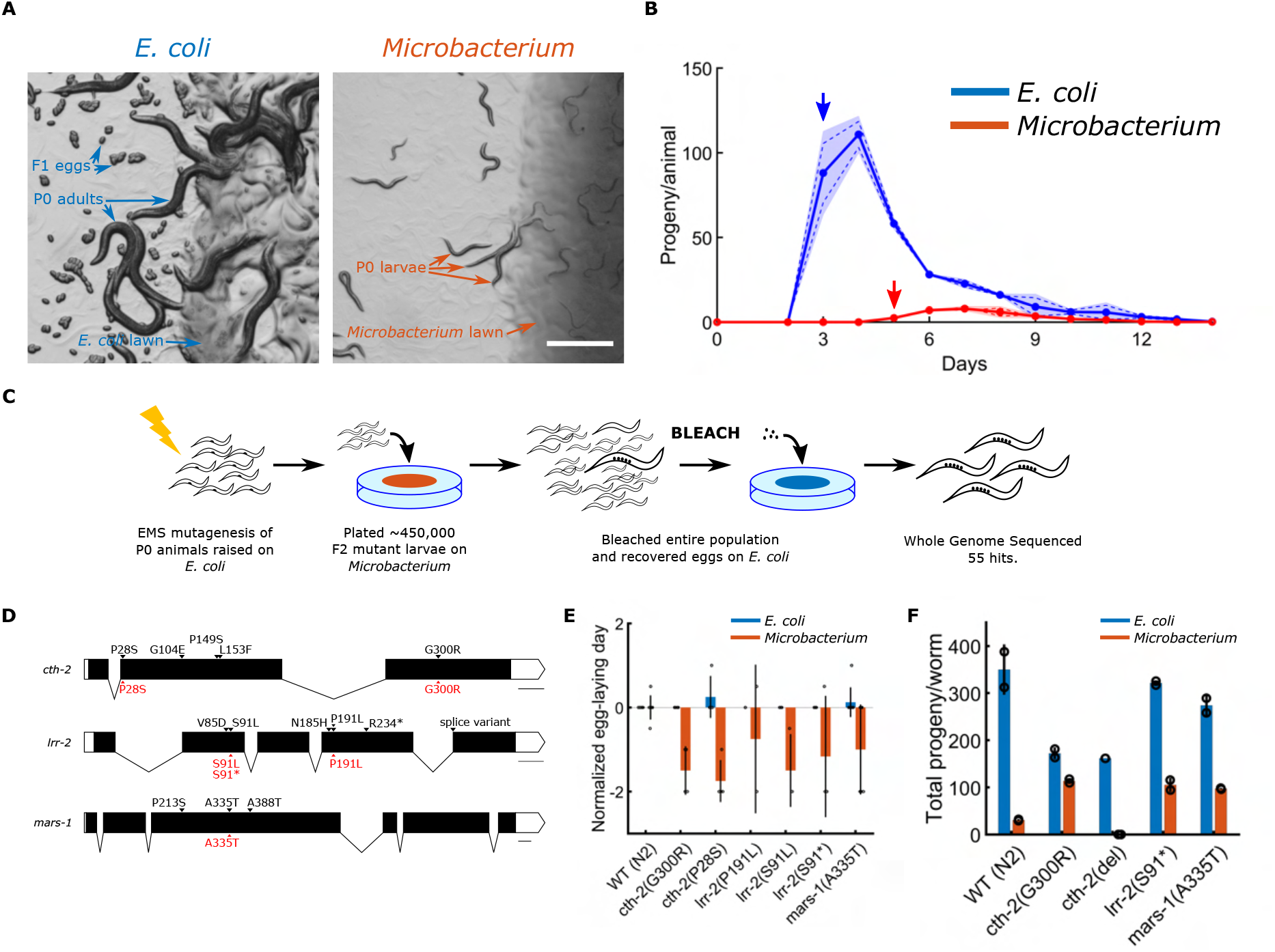
*Microbacterium* JUb115, an exemplary *Actinobacteria* species, causes growth retardation and reduced fertility in *C. ele-gans*. (A) Representative images of *Wild Type (WT) C. elegans (N2)* growing on *E. coli* OP50 (left) or *Microbacterium* JUb115 (right) after 3 days of plating synchronized L1-stage larvae. Scale bar is 500 *µ*m. (B) Time course of progeny laid by *WT*. Arrows illustrates the proxy for growth rate used throughout the manuscript - the first day of egg-laying (3 on *E. coli* and 5 on *Microbacterium* for *WT C. elegans*). (C) Schematic illustration of the forward genetic screen used to isolate *C. elegans* mutants that grow faster on *Microbacterium* than *WT*. (D) Three genes were found with multiple alleles in the screen (black, above gene body diagram). *De novo* CRISPR mutants of some alleles were made (red, below gene body diagram). (E) The first day of egg-laying by CRISPR mutants is shorter than *WT* animals on *Microbacterium* but not on *E. coli*, recapitu-lating the screen phenotype. (F) The mutants rescue fertility of animals on *Microbacterium*.

To discover the molecular mechanism of this delay we performed an unbiased forward genetic screen by chemically mutagenizing *C. elegans* and seeking mutants that grow faster on *Microbacterium* (Figure 1C). We used the first day of egg-laying as a proxy for developmental rate (Figure 1B). *C. elegans* grown on *Microbacterium* takes 5 days to lay eggs, 2 days slower than when fed *E. coli.* We raised mutagenized animals on *E. coli* until the F2 generation, when the alkylating agent induced mutations would become homozygous, and plated axenic cultures of synchronized larvae on *Microbacterium*. When exposed to hypochlorite solution, *C. elegans* tissue dissolves, but its eggs remain unharmed. Thus, after 3 or 4 days of growth we selected for eggs that might have been produced in rare, fast growing mutant F2 animals by treating the animals to hypochlorite solution in the Bleach Screen (Figure 1C, see methods for details). We found no mutants (0 of 200,000 animals tested) that produced eggs in 3 days. We did find mutants with early egg-laying on Day 4 (421 of 460,000 animals). A secondary screen of viability on *E. coli*, and recapitulation of early growth on *Microbacterium* filtered the mutants down to 67, of which 54 were whole-genome sequenced (see Supp. Tab 1 for details).

Each of the sequenced strains contains 100s of point mutations. We identified probable causal lesions by seeking genes with multiple, independent mutant alleles that resulted in amino acid substitutions or other strong effects in the isolated mutants. We identified such mutations in *cth-2* (CystaTHionine gamma-lyase, CTH in humans), which produces cysteine from cystathionine and H_2_S from cysteine; in *mars-1* (Methionyl Amino-acyl tRNA Synthetase, MARS1 in humans), which charges methionine-tRNA with methionine; and in an uncharacterized gene *Y42G9A.3*, which we here name *lrr-2* (Leucine Rich Repeat) due to its homology to LRRC58 in humans^31^ (Fig. 1D).

Using CRISPR we created *de novo* mutants of several probable causal lesions in *WT* animals (Figure 1D, and Supp. Tab. 2). The two alleles *cth-2(P28S)* and *cth-2(G300R)* recapitulated the screen phenotype. However, they are not null alleles. We tested a previously isolated null, *cth-2(W21*)*^13^, and generated a new null allele wherein we deleted a large region of *cth-2* using CRISPR and two guide RNAs, *cth-2(del).* Both of these null alleles caused *C. elegans* to arrest in their larval state and fail to produce a brood when grown on *Microbacterium* (Fig. 1F,5, Supp. Fig. 1).

We hypothesized the *lrr-2* alleles were null alleles because one lesion was a splice variant, which should lead to Nonsense Mediated Decay. Thus, we created an early stop mutant with CRISPR, *lrr-2(S91*),* in addition to *lrr-2(S91L)* and *lrr-2(P191L)* found in the Bleach Screen; all of which recapitulated the screen phenotype of allowing animals to lay eggs earlier than *WT* animals fed *Microbacterium* (Figure 1E). The allele *mars-1(A335T)* that we created with CRISPR also recapitulated the screen phenotype, and is unlikely to be a null allele because *mars-1* is an essential gene. Furthermore, all rescue mutations markedly improved their brood size on *Microbacterium*. Mutants of *lrr-2* and *mars-1* did not suffer reduced brood sizes when fed *E. coli*; however, mutants of *cth-2* did (Figure 1F, Supp. Fig. 1).

### 2. H_2_S gas rescues developmental arrest of animals lacking endogenous cysteine production grown on *Microbacterium*

Cystathionine gamma-lyase, *cth-2*, is conserved throughout the tree of life. Furthermore, the specific amino acid residues that were selected in the Bleach Screen are well conserved across the tree (Figure 2A, Sup. Fig. 2). Despite this conservation, the amino acid substitutions that emerged from the screen are not null alleles of *cth-2*, as null alleles cause larval arrest of *C. elegans* grown on *Microbacterium* (Figure 2C). The main functions of *cth-2* are to metabolize cystathionine to cysteine and cysteine to H_2_S (Figure 2B)^26,32^. To determine how these functions modulate animal development and are affected by the point mutations and null alleles of *cth-2,* we performed a metabolite supplementation screen. Since *cth-2* and *mars-1* are in the sulfur metabolism pathway, we tested a series of metabolites associated with this pathway by supplying them exogenously to *C. elegans* strains fed either *E. coli* or *Microbacterium* (Supp. Fig. 3). We found addition of cysteine or N-acetyl cysteine to the agar plates rescued the larval arrest of *cth-2(del)* on *Microbacterium*, while no other metabolite, including methionine, could (Sup. Fig. 3), suggesting *cth-2(del)* animals lack cysteine when growing on *Microbacterium*. In contrast, several metabolite supplements abrogated the rescue phenotype of *cth-2(G300R)* on *Microbacterium*: methionine, S-adenosyl methionine, homocysteine, cysteine, N-acetyl cysteine, and the CTH inhibitor DL-propargylglycine^33^. This suggested that *cth-2(G300R)* animals grow better on *Microbacterium* because they reduce cysteine levels to an optimal level compared to *WT* animals.

**Figure 2.**
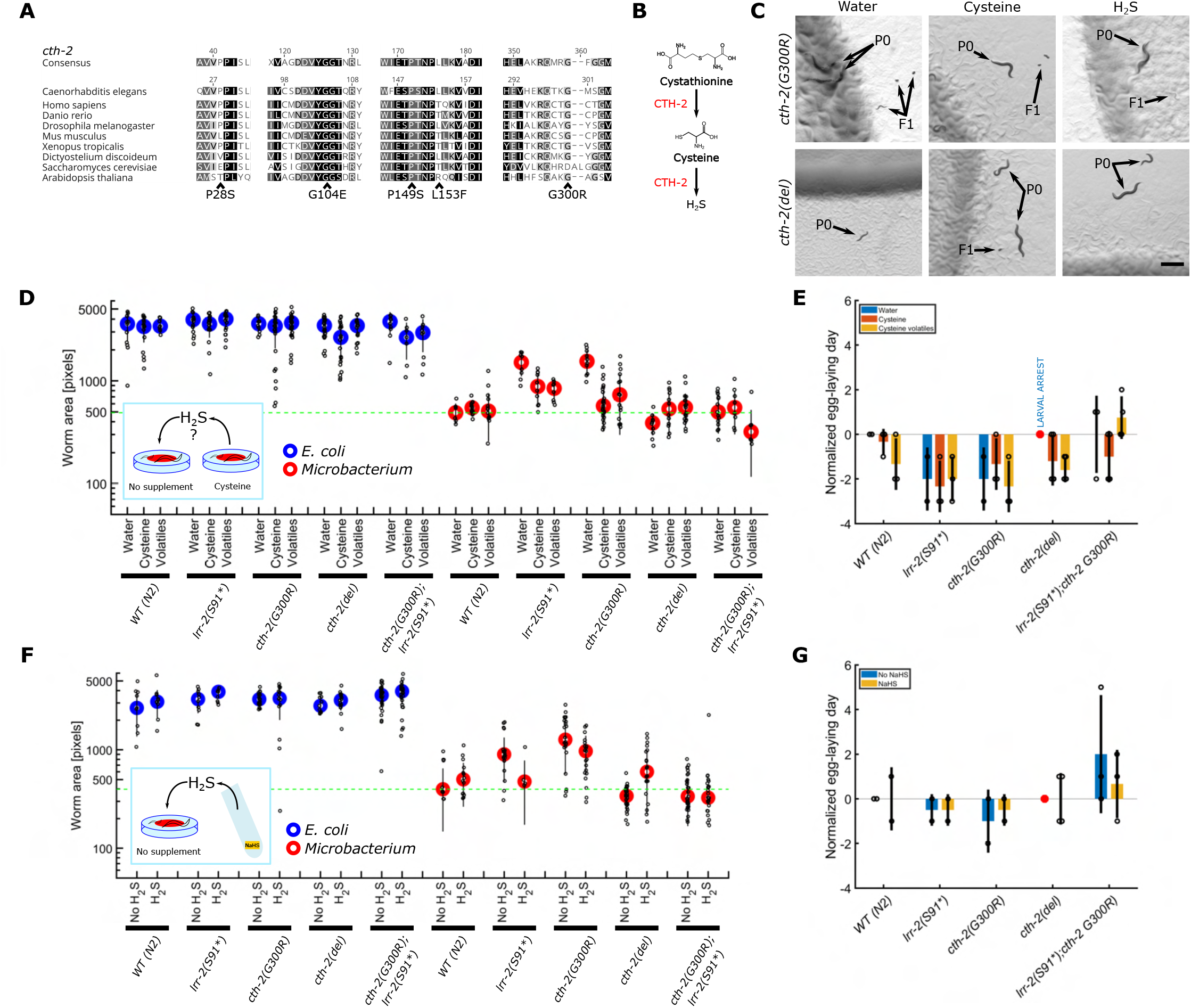
H_2_S is sufficient to rescue animals lacking endogenous cysteine production, but cysteine and H_2_S are deleterious to *lrr-2* and *cth-2* mutants that improve *C. elegans* growth on *Microbacterium*. (A) Residues of *cth-2* identified in bleach screen are well conserved in the tree of life. Alleles identified in the Bleach Screen are indicated below the alignments. (B) Schematic of the functions of *cth-2* : it metabolizes cystathionine to cysteine, and cysteine to H_2_S. (C) Representative images of *C. elegans* strains with *cth-2(G300R)* and *cth-2(del)* alleles on *Microbacterium* with addition of water, cysteine, or exposed to the H_2_S released from cysteine after 5 days of growth. P0 animals and F1 progeny are indicated by arrows. Scale bar is 500 *µ*m. (D) *C. elegans* were grown on plates seeded with either *E. coli* or *Microbacterium* supplemented with 100 *µ*M cysteine or on plates with no supplementation but in the same air-tight container, as shown in the inset schematic. Plates treated with water were sealed in a separate air-tight container as a mock control. Worm lengths were measured 3 days after plating synchronized L1-stage larvae. The data is plotted on a log-scale, each circle represents an individual animal, the blue/red dot represent the median, the error bar is the standard deviation. The dashed line is at the median size of *WT* on *Microbacterium* with control water treatment, as a guide to the eye. (E) The first day of egg laying for the experiments described in (D) on *Microbacterium* was also measured. Each dot represents an independent experiment, bars are mean, error bars standard deviation. Red dot indicates larval arrest. All data is normalized to *WT* on *Microbacterium* with water. (F) *C. elegans* were grown on plates of either *E. coli* or *Microbacterium* placed in an air-tight container with a piece of NaHS in a tube, which releases H_2_S. Data plotted as in (D). (G) First day of egg-laying for the experiments described in (F), and plotted as in (E).

We observed that solutions of cysteine produced a pungent odor, and wondered if the volatiles released by the non-enzymatic degradation of cysteine (H_2_S and NH_3_)^34^ might be sufficient to rescue *cth-2(del)* animals and abrogate the rescue of *cth-2(G300R).* To test this idea, we placed plates that had no supplementation in a sealed container with other plates that were supplemented with cysteine – all seeded with either *E. coli* or *Microbacterium* (see methods for details). We found *cth-2(del)* animals grown on plates seeded with *Microbacterium* and with no supplementation but placed in the same sealed container as the cysteine supplemented plates were rescued as well as those animals on cysteine supplemented plates (Figure 2C-E). This implicated the volatile compounds released from the degradation of cysteine in the rescue of *cth-2(del)*.

To determine whether the causal volatile compound was H_2_S, we placed untreated plates in a sealed container with solid sodium hydrosulfide (NaHS), which hydrolyses spontaneously to gaseous H_2_S. Hydrogen sulfide supplied in this manner rescued *cth-2(del)* as well as cysteine supplementation did (Fig. 2F,G). H_2_S also had a deleterious effect on *cth-2(G300R)* animals similar to cysteine (Figure 2F,G).

*Mycobacterium tuberculosis*, the etiological agent of tuberculosis from the *Actinobacteria* clade, has been found to benefit from host produced H_2_S^23,24^. Thus, we reasoned that the reduced cysteine and H_2_S levels of *cth-2(G300R)* mutants might develop faster because *Microbacterium* pathogenesis relies on *C. elegans* produced H_2_S. We tested this hypothesis by growing *Microbacterium* on plates supplemented with water, cysteine, or methionine; or on plates placed in airtight containers with or without NaHS. Remarkably, we found that cysteine and H_2_S are bacteriostatic but not bactericidal to *Microbacterium*, disproving the hypothesis that *C. elegans* derived H_2_S is beneficial to *Microbacterium* (Figure 3).

**Figure 3:**
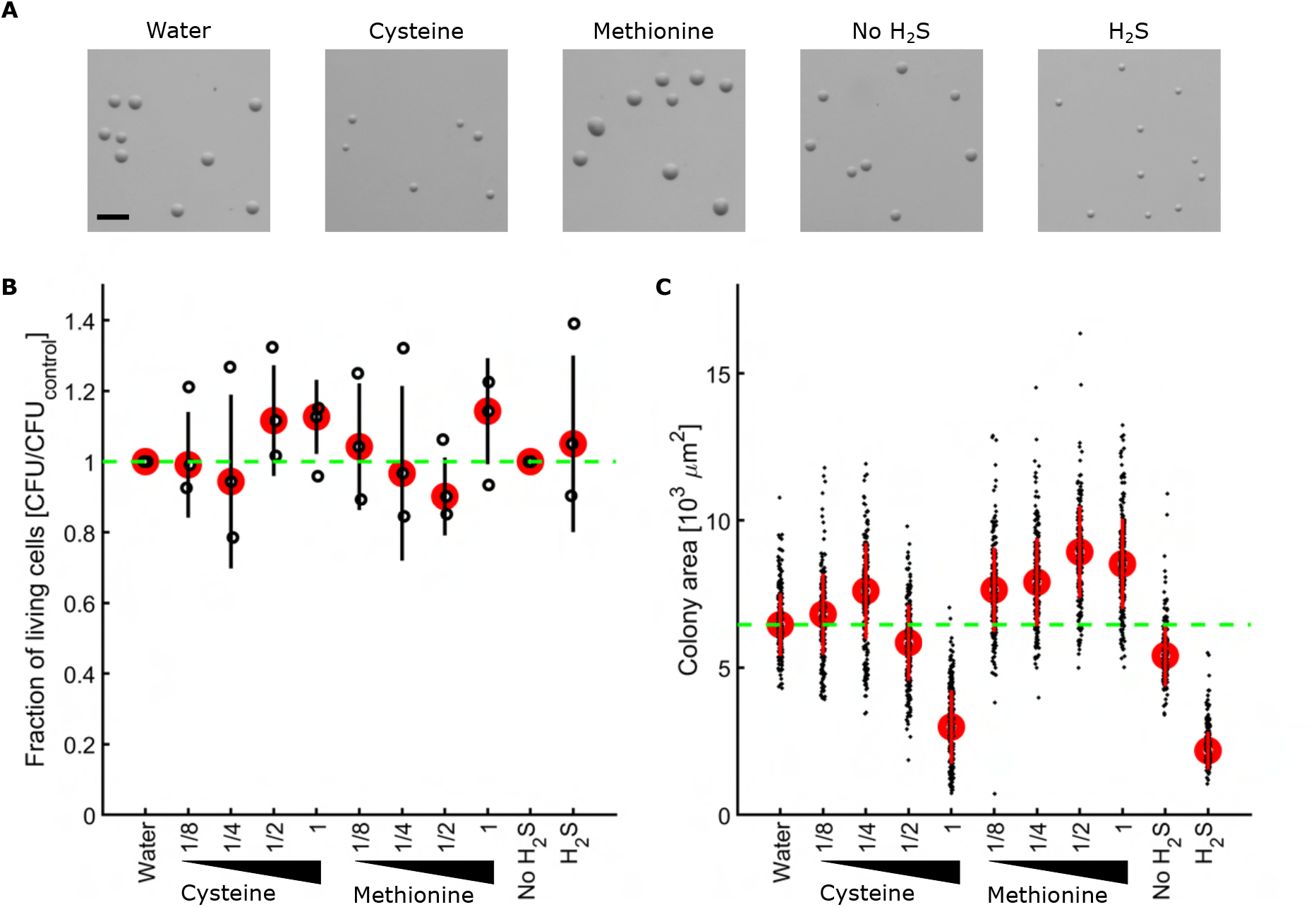
Cysteine and H_2_S are bacteriostatic, not bactericidal, to *Microbacterium*, but methionine is beneficial. *Microbacterium* overnight cultures were diluted and spotted on plates with water, cysteine (dilution series starting at 100 *µ*M), methionine (dilution series starting at 100 *µ*M), or placed in an airtight container with nothing or NaHS. (A) Representative images of colonies for these conditions. Scale bar is 200 *µ*m. (B) Colony forming units were counted and plotted normalized to water for methionine and cysteine, or nothing for H_2_S. Each open circle represents a replicate, red circle the median, error bar the standard deviation. The green dashed line is a guide to the eye at 1. (C) The colony size of each condition in (A) was measured. Each open circle represents a colony, red circle the median, error bar the standard deviation. The green dashed line is a guide to the eye at the median value of colony size on water.

### 3. Uncharacterized cysteine homeostasis regulator acts via cysteine dioxygenase to rescue animal fertility

The uncharacterized gene *lrr-2* is homologous to LRRC58 in humans and is well conserved in animals (Figure 4A). In a high-throughput bait-and-prey experiment with human cells LRRC58 was found to be prey to CDO1, ATG14, and CUL5 baits^35^. CDO1 acts downstream of CTH to oxidize cysteine into cysteinesulfinate. Thus, we reasoned that the LRR-2 protein in *C. elegans* binds CDO-1. Furthermore, CUL5 is a cullin protein, a component of a ubiquitin ligase complex^36,37^ and CDO is known to be poly-ubiquitinated in rats fed cysteine poor diets^38^. Thus, we further hypothesized that LRR-2 may be the recognition domain of the ubiquitin ligase complex and represses CDO-1 by targeting it for ubiquitination (Figure 4B). Fortuitously, a *cdo-1(C85Y)* mutant previously isolated^13^ grows poorly on *Microbacterium* (Figure 4C, Sup. Fig. 1). If *lrr-2* acts via *cdo-1*, *cdo-1* should be epistatic to *lrr-2* and the double of these mutants should behave like *cdo-1*, which is what we observed (Figure 4C).

**Figure 4.**
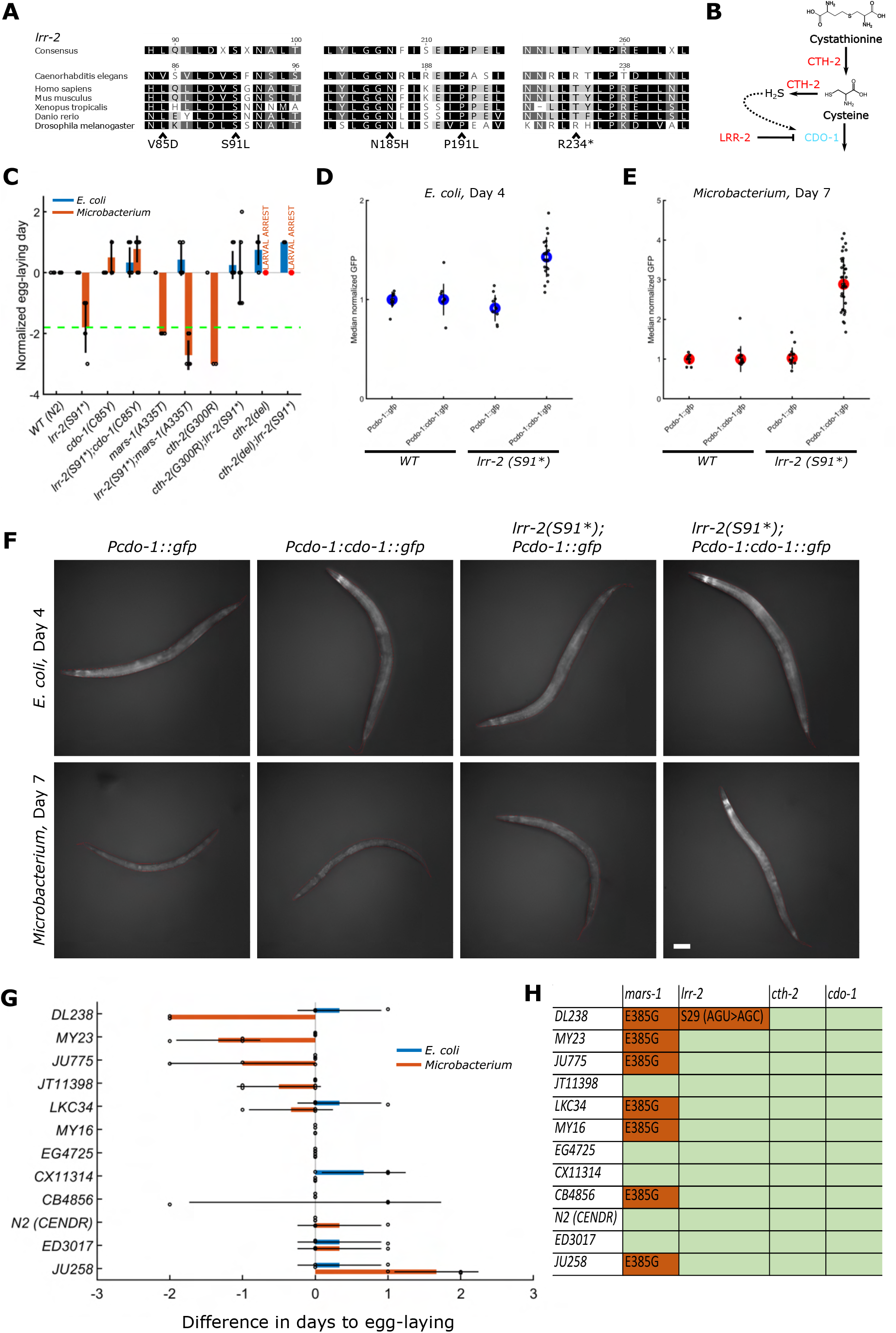
Uncharacterized Leucine Rich Repeat gene, *lrr-2* regulates cysteine levels by post-translationaly regulating *cdo-1*, downstream of *cth-2*. (A) *lrr-2* is homologous to LRRC58 in humans and is conserved across animals. (B) Schematic illustrating the hypothesized interaction of LRR-2 with CDO-1; LRR-2 is a protein that tags CDO-1 for degradation in a ubiquitin-ligase complex, based on the affinity of LRRC58 for CDO in humans. (C) Doubles of *lrr-2* with *cdo-1*, *cth-2(G300R)*, *cth-2(del)*, *mars-1(A335T)* demonstrating that *lrr-2* is epistatic to *cdo-1* and *cth-2*. Each dot is an independent experiment, normalized to *WT* grown on the respective bacteria, bars are mean, error bars are standard deviation. (D,E) Transcriptional and translational reporters crossed into *lrr-2(S91*)* background raised on *E. coli* for 4 days (D) or *Microbacterium* for 7 days (E). Background signal was subtracted and data normalized to the *WT* reporter median: (*T − T_bkg_*)*/W − W_bkg_*), where *T* represents strain plotted and *W* the *WT* reporter. (F) Representative images of animals presented in (D,E). Red outline defines the boundary of the worm. (G) Natural isolates of *C. elegans* were grown on *Microbacterium* JUb115 or *E. coli* OP50 and their days to egg-laying measured. Dots represent individual experiments, bars the mean, lines standard deviation. (H) The wild isolates have mutant alleles of *mars-1* and *lrr-2* but not of *cth-2* and *cdo-1*. The most significant lesion of *mars-1* is shown. *LKC34, JU775, JU258, DL238*, and *CB4856* have additional mutations in *mars-1*.

**Figure 5:**
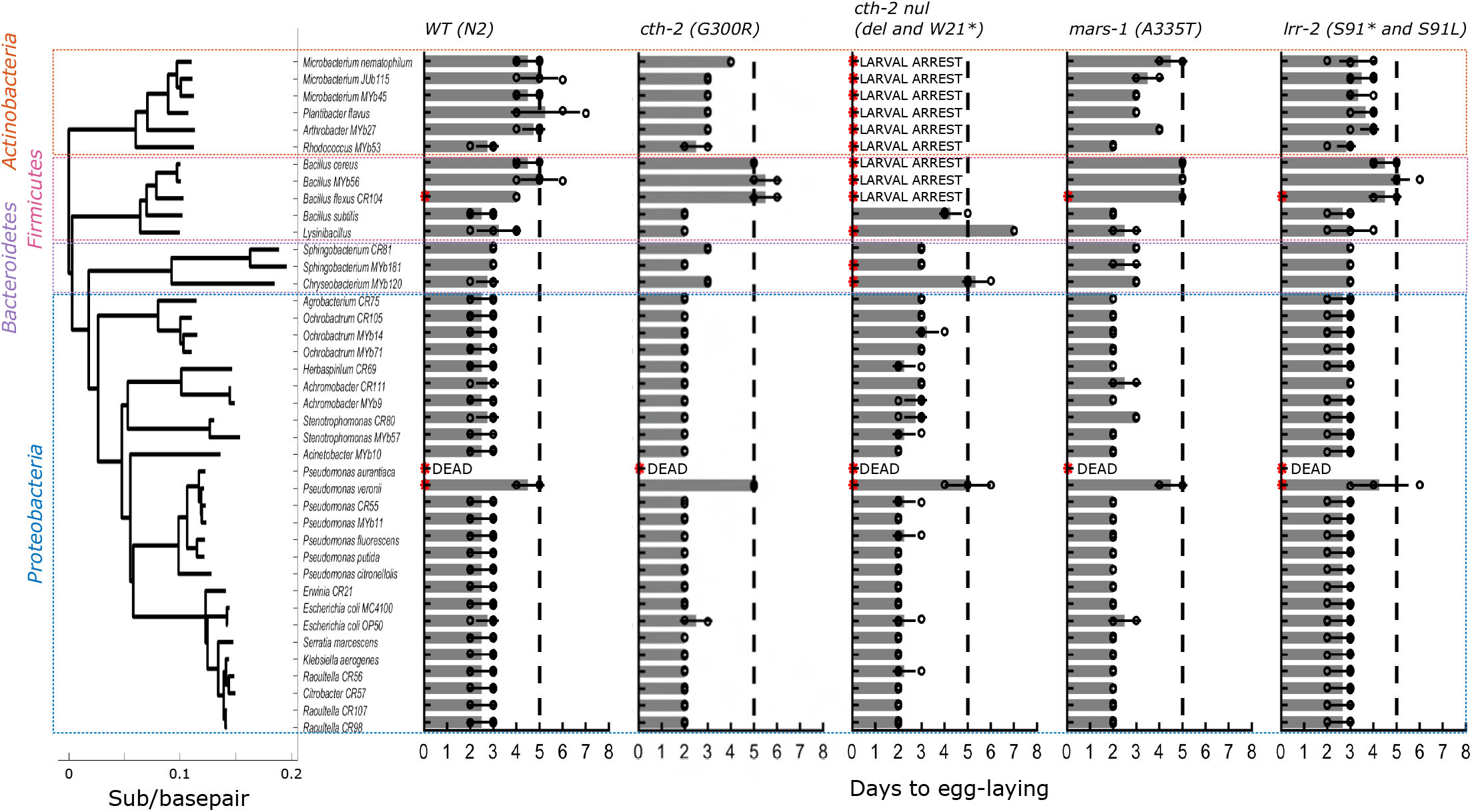
Mutants isolated on *Microbacterium* JUb115 have a similar phenotype on most other *Actinobactera*, but not on *Firmicutes* or *Bacteroidetes*. Phylogeny of bacterial species tested (left). First day to egg-laying for *WT* ; *cth-2(G300R)*; *cth-2(del)* and *cth-2(W21*)*; *mars-1(A335T)*; and *lrr-2(S91*)* and *lrr-2(S91L)* mutants. Each dot represents an independent experiment, bars the mean, error bars the standard deviation; red dots represent instances where eggs were not observed, or a larval arrest phenotype was observed. Dashed line is at the mean days to egg-laying of *WT (N2)* on *Microbacterium JUb115*as a guide to the eye.

We found that addition of cysteine or H_2_S abrogated the rescue phenotype of *lrr-2(S91*)* fed *Microbacterium*, consistent with its inability to degrade CDO-1 and so to reduce cysteine levels (Figure 2D-G). Furthermore, we crossed *lrr-2(S91*)* with *cth-2(G300R)* and *cth-2(del).* Remarkably, the *lrr-2(S91*);cth-2(G300R)* double mutant had a partially penetrant larval arrest phenotype, similar to *cth-2(del)* alone (Figure 4C, Figure 2D-G). Also, *lrr-2(S91*)* was not able to reverse the larval arrest phenotype of *cth-2(del)* animals on *Microbacterium* (Figure 4C), supporting the model that *lrr-2* is a novel member of the sulfur metabolism pathway. In the absence of *lrr-2*, CDO-1 levels increase, thus, cysteine levels should decrease (Figure 4B). We hypothesized the larval arrest phenotype of *lrr-2(S91*);cth-2(G300R)* is due to the two mutant alleles synergistically reducing cysteine levels to those experienced by *cth-2(del)* animals. A prediction of this model is that cysteine supplementation should rescue the slow growth of *lrr-2(S91*);cth-2(G300R)*, which is what we observed (Figure 2D). However, H_2_S causes *lrr-2(S91*);cth-2(G300R)* to grow worse on *Microbacterium* (Figure 2C-F).

We tested this post-translational regulation model by crossing transcriptional (*Pcdo-1::gfp)* and translational (*Pcdo-1::cdo-1::gfp)* reporters of *cdo-1*^39^ into the *lrr-2(S91*)* background. Cysteine and its degradation product hydrogen sulfide (H_2_S) are known to upregulate *cdo-1* in order to maintain cysteine homeostasis^38,39^. Thus, in the absence of *lrr-2*, CDO-1 protein levels should increase, causing a decrease in cysteine levels due to greater degradation, which in turn should cease the upregulation of *cdo-1* transcription. This is what we observe: when *lrr-2(S91*);Pcdo-1::gfp* animals are grown on either *E. coli* or *Microbacterium*, GFP expression is either slightly lower or similar to *WT Pcdo-1::gfp* animals, respectively, but GFP expression in *lrr-2(S91*);Pcdo-1::cdo-1::gfp* animals is markedly increased compared to *WT Pcdo-1::cdo-1::gfp* animals (Figure 4D-F).

### 4. The *C. elegans* sulfur metabolism pathway discovered with *Microbacterium* is ecologically relevant

We found that a diverse set of wild isolates of *C. elegans*^40^ have large variation in their developmental timings on *Microbacterium* JUb115 (Figure 4G). Using the existing genome sequences of these strains^40^ we found that about half the strains bear *mars-1* mutations, but these mutations are uncorrelated with growth on *Microbacterium*. None of these strains are mutants in *cth-2* or *cdo-1*. Remarkably, wild isolate *DL238*, which grows the fastest on *Microbacterium* of the 12 strains tested, bears a mutation in *lrr-2. DL238* has a synonymous mutation at serine 29 with a change in codon from AGU, which is used in *C. elegans* at a frequency of 12.1 per thousand, to AGC which has a usage frequency of 8.4 per thousand^41^. The synonymous mutation might act as a hypomorph of *lrr-2* by causing lower translation efficiency. Thus, the specific modifications to the *C. elegans* sulfur metabolism pathway found here may be under selection in the wild.

We raised *WT C. elegans* on a panel of diverse bacterial species from its natural environment as well as type strains from the various clades^7,17,42^ (Fig. 5). Three of the four dominant clades of bacteria in the *C. elegans* natural environment, *Actinobacteria, Firmicutes,* and *Bacteroidetes*, all caused *C. elegans* to develop at a slower rate compared to growth on most members of the *Proteobacteria* clade. To test whether the *C. elegans* sulfur metabolism pathway discovered with *Microbacterium* JUb115 is specific to *Actinobacteria*, we assayed the growth of various mutants on the panel of bacteria. We found that *lrr-2(S91*), lrr-2(S91L), mars-1(A335T),* and *cth-2(G300R)* all caused improved growth on *Actinobacteria*, but no *Firmicutes* nor *Bacteroidetes*. Furthermore, null alleles of *cth-2* caused larval arrest on *Actinobacteria*, and retarded growth on *Firmicutes, Bacteroidetes*, and some *Proteobacteria.* Thus, the specific modifications to the sulfur metabolism pathway discovered with *Microbacterium* JUb115 generalize specifically to the *Actinobacteria* clade.

## Discussion

In this work we discovered the role of hydrogen sulfide and cysteine in mediating the interaction of *C. elegans* with *Actinobacteria*. We began with an unbiased forward genetic screen seeking mutant animals that would escape the growth retardation induced by *Microbacterium*. We demonstrated that the three genes identified are all involved in sulfur metabolism. We found alleles of *cth-2/CTH* that caused developmental arrest on *Microbacterium*, which can be rescued by H_2_S. We also identified a novel regulator of cysteine levels, *lrr-2/LRRC58*. Our discoveries made with *Microbacterium* generalize to the clade of *Actinobacteria* and are likely relevant to wild *C. elegans* populations because lesions in *lrr-2* were detected in a wild isolate of *C. elegans* that can grow well on *Microbacterium*.

Synthesizing our data, we present a conceptual model of how and why H_2_S and the other metabolites in our work mediate animal-microbe interactions. While *Actinobacteria* are not the preferred diet of *C. elegans*, they are evidently nutritionally complete because they support the robust growth of *C. elegans* mutants. In the wild, *C. elegans* might produce H_2_S from its trans-sulfuration pathway to inhibit the growth of *Actinobacteria* to enable its preferred diet, *Proteobacteria*, to thrive. However, when faced with the artificial scenario of a monoculture *Actinobacteria* diet, its evolved response backfires and causes its sole source of nutrition to be diminished, thus causing developmental retardation of *WT C. elegans* on *Actinobacteria*.

The striking rescue of *cth-2(del)* larval arrest on *Microbacterium* by H_2_S is opposite to the deleterious effect of H_2_S on *cth-2(G300R)* and *lrr-2(S91*)*. The *cth-2(null)* animals might be made so ill by the lack of endogenous cysteine production that they require their diet to be deleteriously affected by H_2_S for consumption. Another, more likely, possibility is that H_2_S is metabolized by *Microbacterium* into cysteine by a homologue of *Mycobacterium tuberculosis* O-Acetylserine Sulfhydrylase (CysK1)^25^. Thus, *cth-2(null)* animals may be getting the required cysteine levels because, under H_2_S exposure, *Microbacterium* has an increased abundance of cysteine. The animals may also directly convert the exogenous H_2_S to cysteine using CYsteine Synthase Like genes (*cysl-1/2*)^14^.

Our discovery of the function of *lrr-2/LRRC58* might be combined with the observation that H_2_S is beneficial to *Mycobacterium tuberculosis* for therapeutic purposes. It would be intriguing to find that *LRRC58* mutants are better able to survive against *Mtb*, similar to *CTH* and *CBS* mutants^23,24^. If this were the case, the leucine-rich repeat of *LRRC58* might be a convenient drug target to reduce the virulence of *Mtb* in patients, where it would act to reduce the H_2_S available to the infecting bacteria.

Organisms must maintain sulfur homeostasis despite diverse ecological challenges. Our work sheds light on the delicate balance that has evolved in an animal-microbial ecology found on rotting vegetation and demonstrates the surprises that await us as we explore these ecologies in ever greater detail.

## Acknowledgements

OP thanks Giulia Arsuffi for technical assistance in image analysis as well as many fruitful discussions.

## Author contributions

OP and GR designed experiments; wrote and edited the manuscript. OP and PB performed experiments.

## Declaration of interests

The authors declare no conflicts of interest.

## Methods

### Strains and animal culture

*C. elegans* strains were propagated using standard procedures on a diet of *E. coli* OP50^43^, being careful to avoid starvation before any assays. All *C. elegans* experiments were carried out at 20°C. The strains used are listed in Sup. Table 1.

### Bacterial culture

*E. coli* OP50 and *Microbacterium* JUb115 were streaked out on Luria-Bertani (LB) agar plates. Single colonies were picked into liquid LB and grown with shaking overnight at 37°C. 100 uL of the overnight cultures were plated on Nematode Growth Medium (NGM) plates without antibiotics. The plates were incubated overnight at 20°C and then stored at 4°C until they were used (typically within 7 days). For propagation of animals *E. coli* OP50 was seeded on NGM plates with streptomycin and nystatin; these plates were occasionally also used for animal growth assays.

### Bleach treatment

To isolate eggs from adult animals, they were first washed into M9 buffer and allowed to settle. The solution volume was brought to 5 mL by aspiration. To this 2 mL of stock bleach solution (1 mL of 5 M NaOH, 2 mL of 6% hypochlorite solution) was added and incubated with vortexing for 4 minutes. The solution was centrifuged at 3 krpm for 20s, the supernatant discarded, and the precipitate washed 3x in 10 mL M9 buffer. The resulting eggs were either synchronized in 3 mL of M9 buffer or on unseeded NGM agar plates overnight at 20°C.

### Bleach screen

*WT (N2)* animals were propagated on Nematode Growth Medium (NGM) plates seeded with *E. coli* OP50. Synchronized L1 larval stage animals were plated on OP50 and designated the Parental generation (P0). After 2 days of growth, L4 larval stage animals were washed in M9 buffer and incubated with ethyl methanesulfonate (EMS) for 4 hours. Animals were washed several times in M9 and then plated on fresh OP50 plates to recover. F1 eggs were isolated with bleach treatment and F1 synchronized L1 larvae plated on OP50. After 3 days, F2 eggs were isolated with bleach treatment. F2 synchronized L1s were plated on NGM plates seeded with *Microbacterium*. In the first iteration of the screen, half the *Microbacterium* plates were treated with bleach 3 days after plating the animals, and the other half 4 days after. As the first round of screening produced no mutants on Day 3, the second round of screening only involved bleaching the *Microbacterium* plates on Day 4 (see Sup. Table 1). The F3 hits from *Microbacterium* plates were then plated on OP50 plates to recover. Over subsequent days, before the F3 animals became gravid, animals were singled out onto fresh OP50 plates. These constituted the mutants found in the Bleach Screen.

### Secondary screen

Strains were propagated on *E. coli* OP50. L4 animals were picked to fresh OP50 plates. After 1-2 days, 5 gravid adults were picked into a 10 uL drop of bleach solution (15 mL bleach, 5 mL 5 M NaOH, 30 mL water) on plates seeded with either *E. coli* or *Microbacterium*. Plates were observed every day for the first day of egg-laying.

### Identifying candidate causal genes

Candidate causal genes in strains that passed the secondary screen were identified using a previously published approach^44^. Briefly, genomic DNA from the strains was prepared using a Gentra Puregene Tissue Kit (Qiagen), and libraries prepared using standard reagents (New England Biolabs). DNA libraries were sequenced on an Illumina NextSeq 2000 and reads were analyzed using a standard workflow followed by a custom MATLAB script.

### Constructing CRISPR strains of candidate mutations

The CRISPR RNA guides were designed with the aid of Geneious and homology repair templates were designed following the principals in Arribere *et al.*^45^. CRISPR injection mixtures were prepared following the protocol of Ghanta and Melo^46^.

### Egg-laying assay

*C. elegans* strains were cultured on *E. coli* OP50 and synchronized larvae were harvested from adults by bleaching and hatching eggs on agar plates without bacterial food (Metabolite Screen, Microbiome Screen). Alternatively, 5 adults (1-2 days old) were dropped in 10 uL of bleach solution (15 mL bleach, 5 mL 5 M NaOH, 30 mL water). Starting on Day 2 after plating animals the plates were inspected manually for egg laying and the first day an egg was observed is reported.

### Fertility assay

*C. elegans* strains were cultured on *E. coli* OP50 and synchronized larvae harvested from adults by bleaching and hatching eggs on agar plates without bacterial food. Between 20 and 40 larvae of each strain were plated on NGM agar plates seeded with either *E. coli* OP50 or *Microbacterium* JUb115. After 2 days of growth on *E. coli*, 12 L4 stage larvae were moved to fresh *E. coli* plates. The next day these P0 animals were moved to fresh *E. coli* plates and the F1 progeny (eggs, hatched larvae, unfertilized oocytes) were counted on the plates the P0 were moved from. This was repeated for up to 14 days or until all P0 adults had died. The animals on *Microbacterium* were treated similarly. However, *lrr-2(S91*), cth-2(G300R), mars-1(A335T)* mature larvae were transferred for the first time on Day 3, not Day 2; between 10 and 12 mature Wild Type *(N2)* larvae on Day 4; between 8 and 12 mature *cdo-1(C85Y)* larvae on Day 5; and no larvae of *cth-2(del)* were transferred to fresh plates as none matured.

### GFP measurements

Synchronized larvae were plated on *E. coli* OP50 or *Microbacterium* JUb115 seeded 9 cm NGM plates. After 4 days of growth on *E. coli*, single adults were picked into a 1-2 uL drop of 1.5 M sodium azide placed on an unseeded, antibiotic free NGM plate. The paralyzed animals were imaged in a Zeiss AxioZoom V16 microscope at 112x magnification with 2 s of illumination in the GFP channel and in brightfield. The same treatment was done for animals on *Microbacterium* after 7 days of growth. The images of animals were traced manually. The area of the image not occupied by the animal was averaged for background signal, which was subtracted from the average pixel intensity in the GFP channel.

### Metabolite screen

Each well of a 24-well plate with antibiotic free NGM was seeded with either *E. coli* OP50 or *Microbacterium* JUb115, and incubated overnight at 20°C. These wells were then treated with either water or 100 uL of 100 mM stock solutions of the metabolites (6 mM for S-adenosyl methionine, 16 mM pyridoxal 5’-phosphate hydrate, and 45 mM for DL-propargylglycine). Synchronized *C. elegans* were plated after the metabolite solutions had dried.

### Cysteine volatiles and H_S_S experiments

To test the effect of cysteine volatiles, synchronized larvae were plated on either *E. coli* OP50 or *Microbacterium* JUb115 seeded plates with either 100 uL of water, 100 uL of 100 mM cysteine, or no treatment. All the water treated plates were placed in one ziplock bag, while all the cysteine or untreated plates were placed in another ziplock bag. To test the effect of H_2_S directly, untreated plates were placed in two ziplock bags. In one bag, a tube with a 20 to 40 mg flake of NaHS was placed. Egg laying was assessed as described above, and length of animals measured on Day 3 as described below.

### *C. elegans* size measurements

Animals were imaged using a Zeiss AxioZoom V16 microscope at 15x magnification in brightfield. The entire plate was imaged by tiling. Outlines of the P0 animals were manually traced in ImageJ, using a script generated with the aid of ChatGPT. The size of the worm is reported as the projected area of the worm in pixels.

### Colony forming units and colony size assay

Unseeded, antibiotic free, 90 mm NGM plates were treated with 225 uL of water, 100 mM cysteine, 50 mM cysteine, 25 mM cysteine, 12.5 mM cysteine, 100 mM methionine, 50 mM methionine, 25 mM methionine, 12.5 mM methionine, or left untreated and incubated at room temperature overnight (∼24°C). Overnight cultures of *Microbacterium* JUb115 were grown in LB at 37°C shaking incubators and were started from single colonies. The overnight cultures were diluted using LB in a 10-fold dilution series, and 5 uL spots of 10^5^ to 10^8^ dilutions were plated on the plates in triplicate. The untreated plates were either placed in a ziplock bag with nothing or a tube with a 62 mg flake of NaHS in it. The plates were incubated at 37°C. The spots of *Microbacterium* were imaged with an Zeiss AxioZoom V16 microscope after 2 days at 15x magnification in brightfield. The centers of the colonies were manually located and used in combination with an image analysis algorithm (created with the aid of ChatGPT) to extract colony size.

### Alignment of amino acid sequences

Amino acid sequences of homologous genes were obtained by NCBI blastp searches. The sequences were aligned in Geneious using Clustal Omega.

### Microbiome egg-laying screen

The collection of bacteria was streaked on LB plates from glycerol stocks and incubated for 1 to 3 days at 37°C. Single colonies were picked into LB and grown overnight at 30°C with shaking. 25 uL of the overnight cultures was plated on antibiotic free, unseeded, 24-well NGM plates. The plates were incubated overnight at 20°C and used immediately, or first stored at 4°C. We note that *Microbacterium* JUb115 is a distinct species from the well-studied *Microbacterium nematophilum,* as it does not cause the Deformed Anal Region (dar) infection phenotype^7^.

### Phylogeny construction of microbes

All bacterial species were 16s sequenced. The identities of the bacteria were found using blastn in Geneious. The appropriate 16s sequences were processed in Geneious with the Tamura-Nei Genetic Distance Model and Neighbor-joining Tree Build Method with *Saccharolobus solfataricus* as the outgroup.

---

**Supplementary Table 1:** Summary of animals assayed in Bleach Screen and subsequently sequenced.

| Quantity | Round 1 |  | Round 2 | Total |
| --- | --- | --- | --- | --- |
| Days post plating F2 mutant L1 larvae on *Microbacterium* | Day 3 | Day 4 | Day 4 | Day 4 |
| F2 mutant larvae plated | 230,000 | 200,000 | 260,000 | 460,000 |
| F3 mutant larvae that survived bleach treatment (% of animals plated) | 0 (0) | 44 (0.02) | 377 (0.14) | 421 (0.09) |
| Secondary screen hits (% of hits from primary screen) |  | 11 (25) | 56 (15) | 67 (16) |
| Mutants sequenced (% of secondary screen hits) |  | 10 (91) | 45 (80) | 54 (81) |
| Causal mutation identified (% of sequenced) |  | 2 (20) | 34 (76) | 36 (67) |
| WT (N2) control F2 larvae plated | 20,000 | 10,000 | 10,000 | 20,000 |
| WT F2 larvae that survived bleach treatment (% of animals plated) | 0 (0) | 0 (0) | 1 (0.01) | 1 (0.005) |
| Secondary screen hits (% of hits from primary screen) |  |  | 1 (100) | 1 (100) |
| Spontaneous mutants sequenced (% of secondary screen hits) |  |  | 1 (100) | 1 (100) |
| Causal mutation identified (% of sequenced) |  |  | 0 (0) | 0 (0) |

**Supplementary Figure 1.**
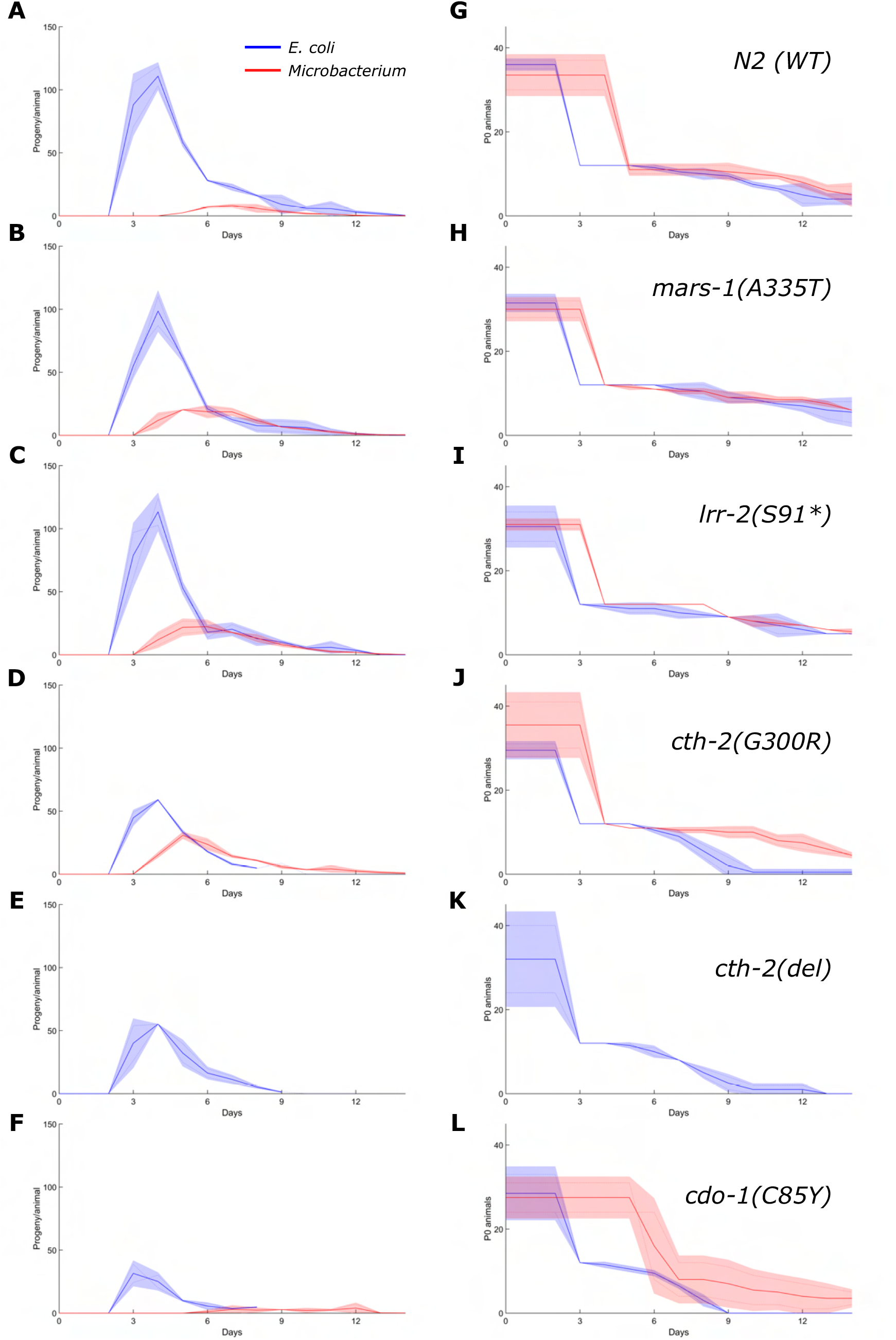
The rescue mutants identified in the bleach screen cause animals to lay eggs earlier and more abundantly over several days. (A-F) Number of eggs laid per P0 worm on given day. Bold line is the mean, shaded area the standard deviation, light lines the individual biological replicates (each condition performed in duplicate). (G-L) Number of P0 worms assayed on given days. For animals grown on *E. coli* 12 L4 worms were transferred to a fresh plate on Day 2 and followed until they died or the time course finished. For animals grown on *Microbacterium*, *mars-1(A335T)*, *lrr-2(S91*)*, *cth-2(G300R)* 12 late stage larvae or young adults were transferred on Day 3; *WT* animals were transferred on Day 5, *cdo-1(C85Y)* animals on Day 5 to 7; *cth-2(del)* animals were never transferred.

**Supplementary Figure 2:**
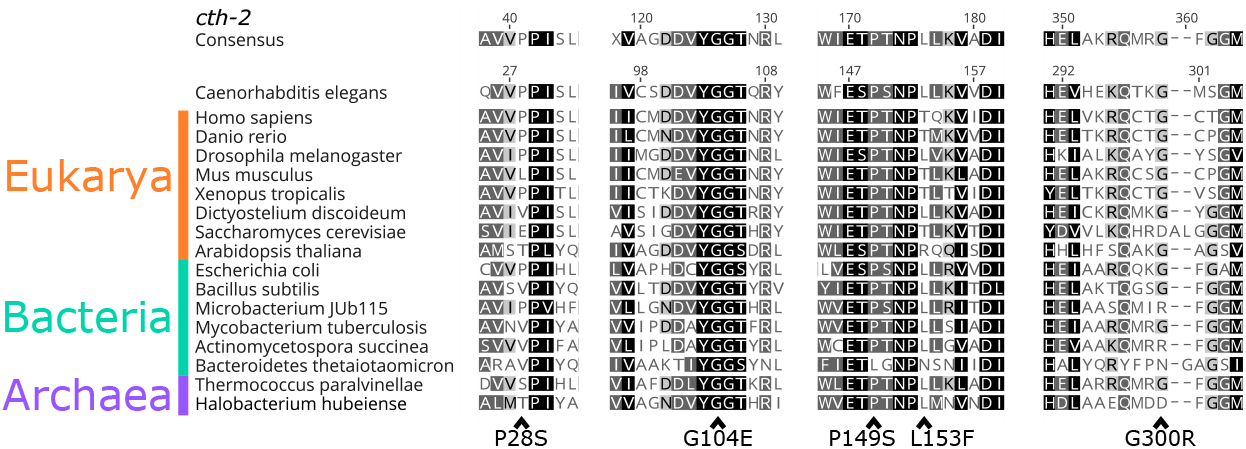
The residues of *cth-2* found in the screen are conserved in the tree of life. Alleles identified in the screen are indicated below the alignments.

**Supplementary Figure 3:**
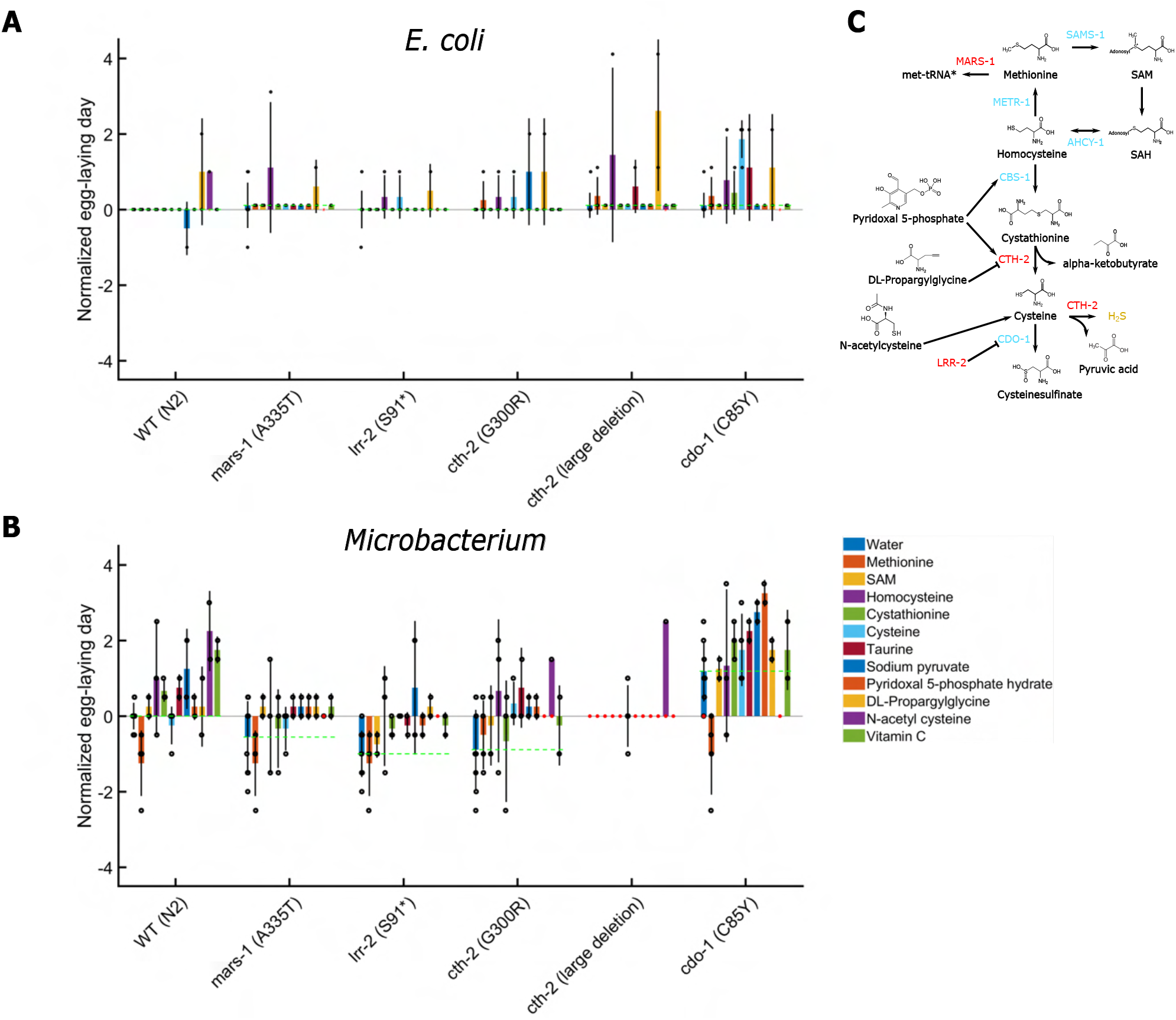
Metabolite supplementation screen reveals cysteine can rescue developmental arrest of *cth-2(del)* animals fed *Mi-crobacterium*. Each dot represents an independent experiment of *C. elegans* strains grown on (A) *E. coli* OP50 and (B) *Microbacterium* JUb115 with ex-ogenous application of metabolites. All experiments done in at least duplicate. The data is normalized by subtracting the mean of *WT C. elegans* grown on *E. coli* (A) or *Microbacterium* (B) supplemented with water (mock control). The dashed line represents the mean of each strain on the respective bacteria with water added. A single egg was occasionally observed to be laid by *cth-2(del)* animals 12 days after plating L1 larvae on *Microbacterium*. However, the P0 animals were still larval sized at this time, thus these events have been classified as larval arrest in the figure. Red stars represent either larval arrest or absence of egg-laying observed. (C) Schematic illustrating the trans-sulfuration (homocysteine to cysteine) pathway and the methionine/S-adenosyl methionine cycle with metabolites used in the screen.

**Supplementary Table 2:**
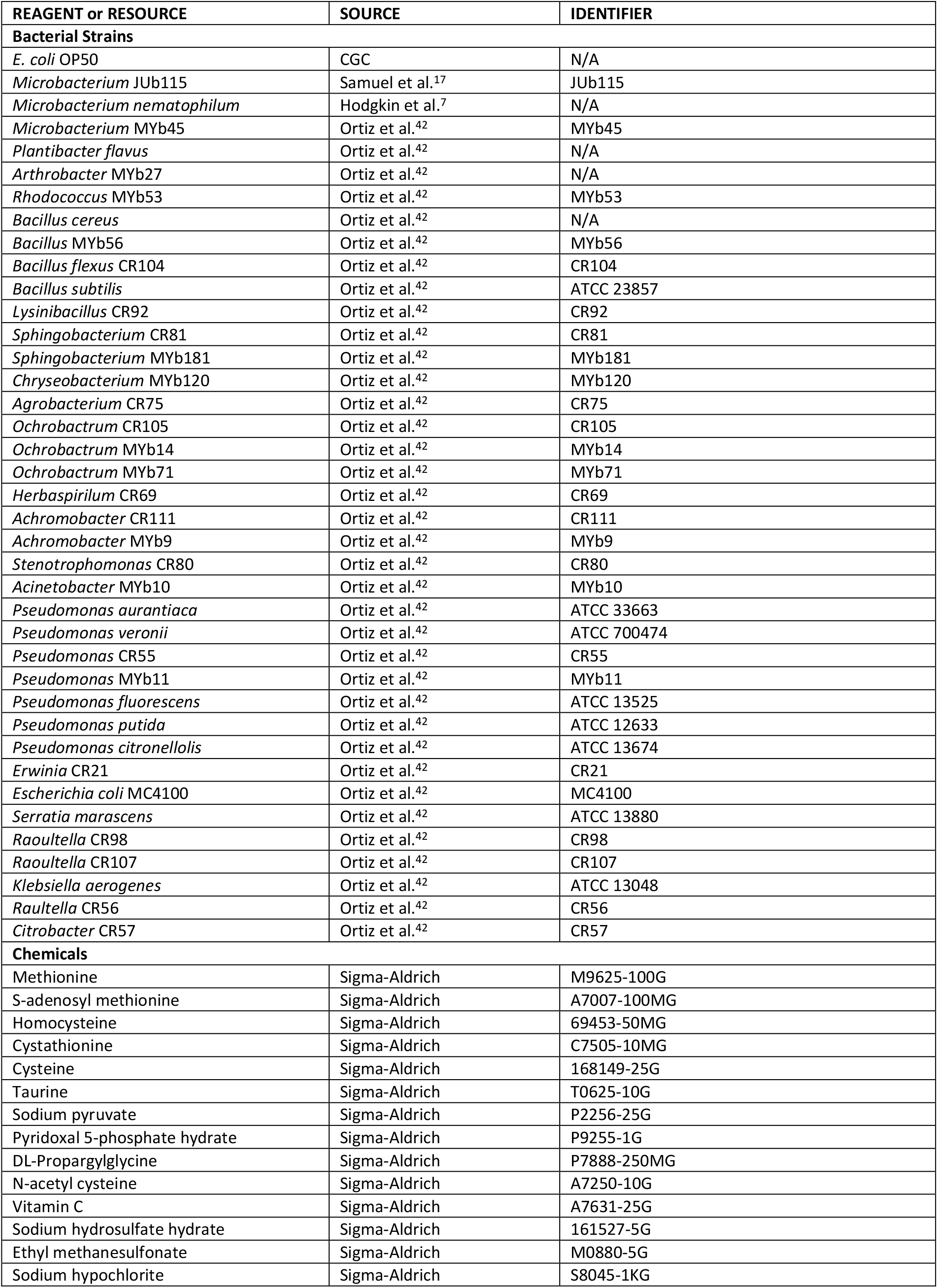
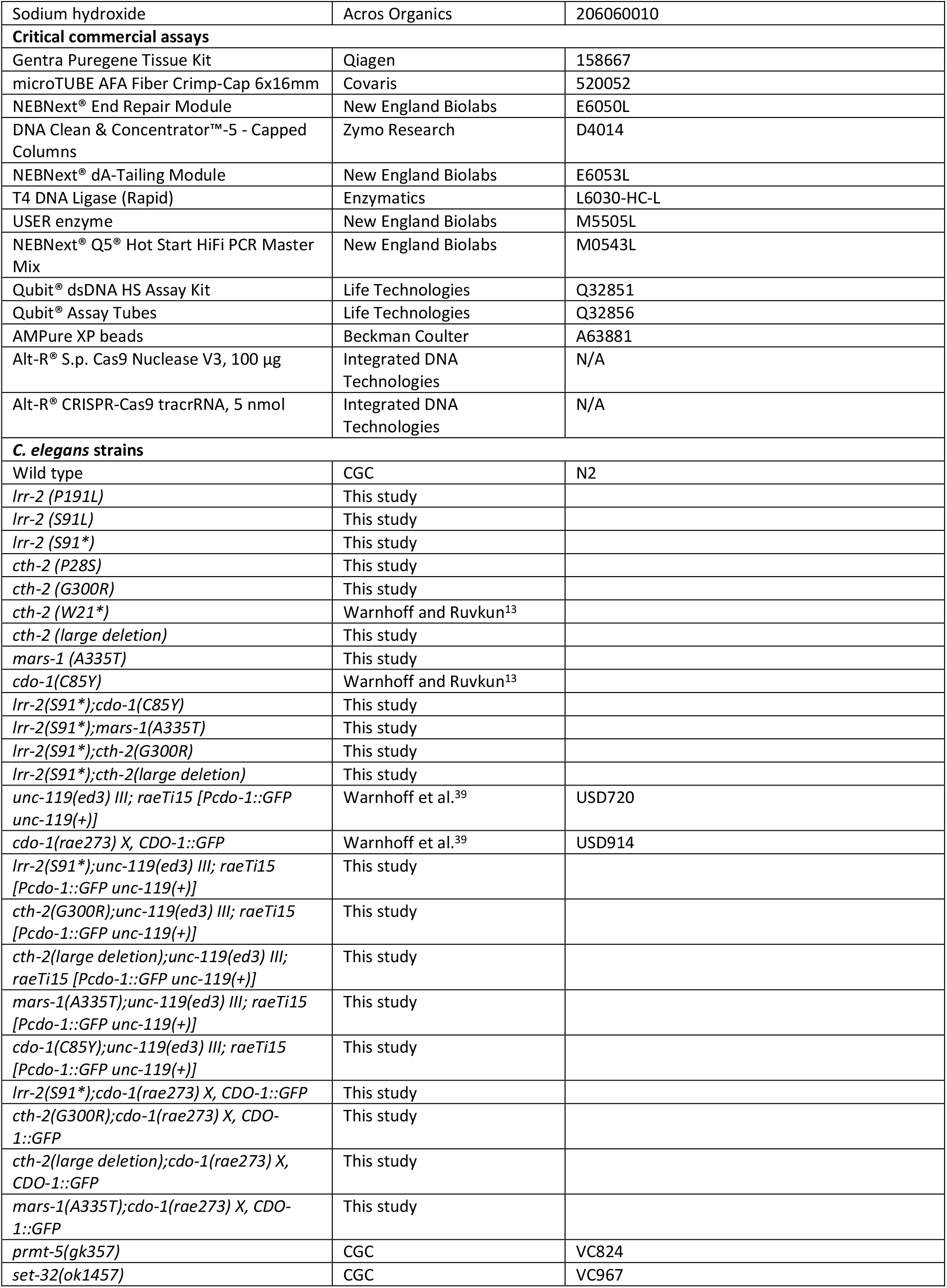
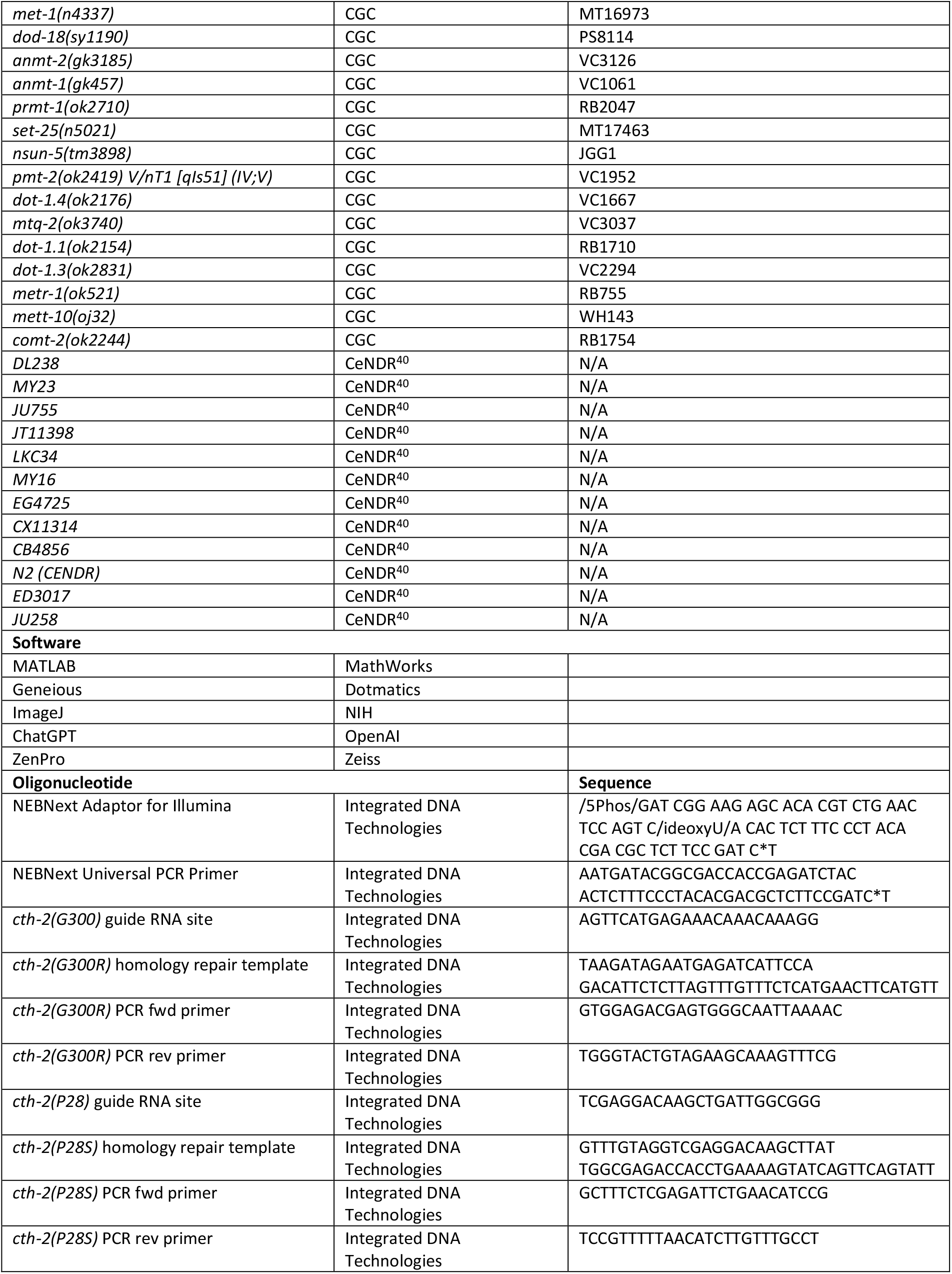
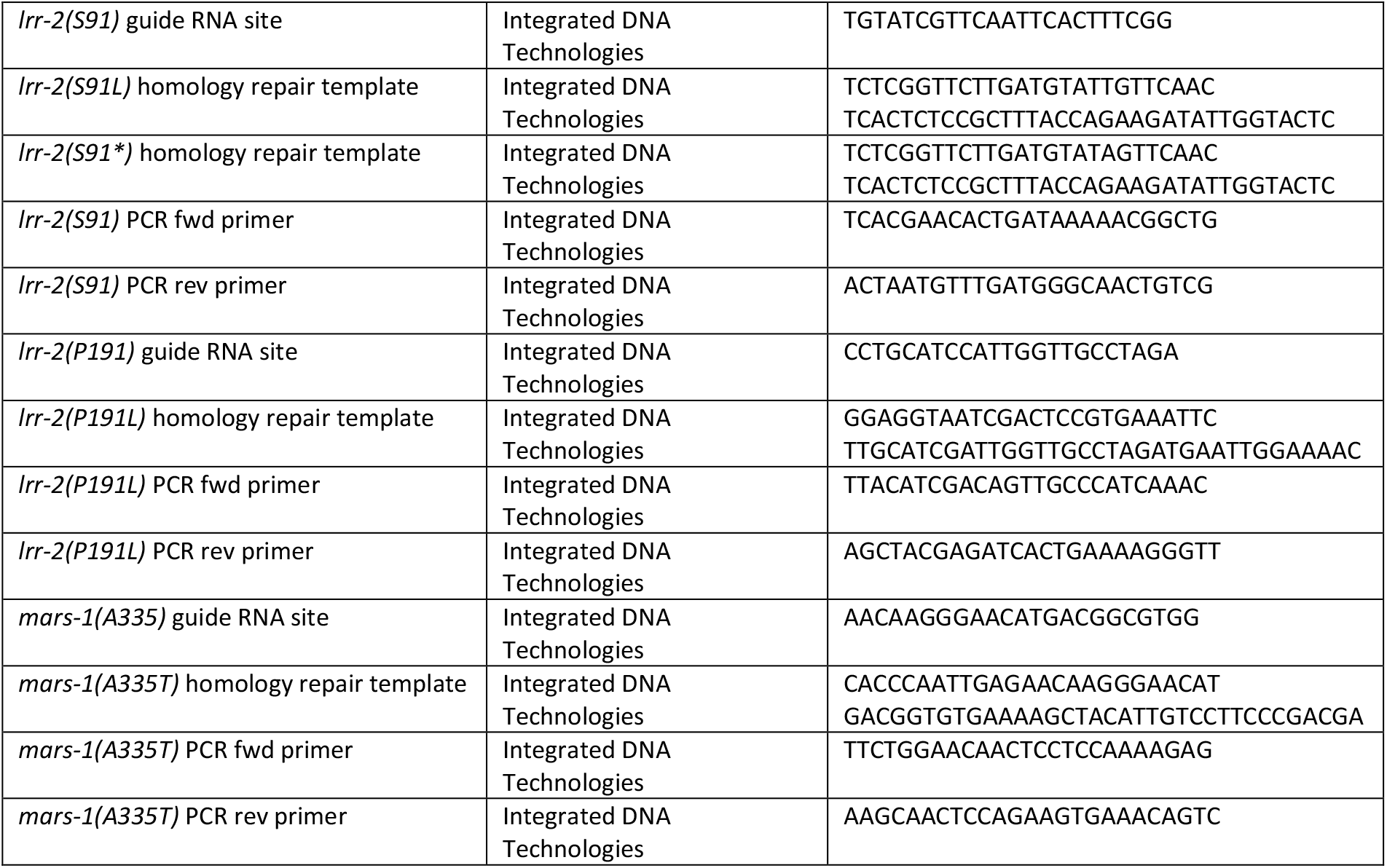
List of reagents and resources.

## Notes

### Competing Interest Statement

The authors have declared no competing interest.

